# A bivalent Adenovirus-Vectored Vaccine induces a robust humoral response, but does not protect cynomolgus macaques against a lethal challenge with Sudan virus

**DOI:** 10.1101/2023.10.20.563337

**Authors:** Sarah van Tol, Paige Fletcher, Friederike Feldmann, Reshma K. Mukesh, Julia R. Port, Shane Gallogly, Jonathan E. Schulz, Joseph F. Rhoderick, Rebecca Makinson, Aaron Carmody, Lara Myers, Jamie Lovaglio, Brian J. Smith, Atsushi Okumura, Carl Shaia, Greg Saturday, Andrea Marzi, Teresa Lambe, Vincent J. Munster, Neeltje van Doremalen

## Abstract

The most recent Sudan virus (SUDV) outbreak in Uganda was first detected in September 2022 and resulted in 164 laboratory-confirmed cases and 77 deaths. Currently, there are no approved vaccines or therapeutics against SUDV. In the current study, we investigated the protective efficacy of ChAdOx1-biEBOV in cynomolgus macaques using a prime or a prime-boost regimen. ChAdOx1-biEBOV is a replication-deficient simian adenovirus vector encoding SUDV and Ebola virus (EBOV) glycoproteins (GPs) at the E1 and E4 loci, respectively. Intramuscular vaccination induced SUDV and EBOV GP-specific IgG responses and neutralizing antibodies. Upon challenge with SUDV, vaccinated animals, regardless of vaccination scheme, showed signs of disease like those observed in control animals, and no difference in survival outcomes were measured among all three groups. Viral load in blood samples and in tissue samples obtained after necropsy were not significantly different between groups. Overall, this study highlights the importance of evaluating vaccines in multiple animal models, including nonhuman primates, and demonstrates the importance of understanding protective efficacy in both animal models and human hosts.

## Background

Of the six viruses described in the genus *Orthoebolavirus (1)*, four are known to cause Ebolavirus disease in humans, namely Ebola virus (EBOV), Sudan virus (SUDV), Bundibugyo virus (BDBV), and Taï Forest virus (TAFV). Although EBOV is responsible for most outbreaks (*2*), SUDV has caused eight outbreaks in Sudan and Uganda since first discovered in 1976 (*3*–*9*). SUDV disease (SVD) has been documented in nearly 1,000 human cases and has a combined case fatality rate (CFR) of 50% (*3*–*9*). The most recent SUDV outbreak was first detected in Uganda’s Mubende district in September 2022 (*9*, *10*). During the months-long outbreak, there were 164 laboratory-confirmed cases and 77 deaths (CFR 47%). The severity of SVD and the increasing potential for wider spread of SUDV outbreaks warrant the development of medical countermeasures.

Current licensed vaccines and therapeutics against EBOV exist, but the cross-protection against SVD in humans has not been demonstrated. The United States Food and Drug Administration (FDA)-approved EBOV vaccine, Ervebo®, utilizes a recombinant vesicular stomatitis virus (rVSV) backbone expressing EBOV Kikwit’s glycoprotein (GP) (*11*). In the pre-clinical stage, Ervebo® protects against lethal EBOV challenge but not SVD in cynomolgus macaques (*12*). A second EBOV vaccine approved by the European Medical Agency utilizes a prime-boost approach of replication-incompetent adenovirus (Ad26) encoding the EBOV Mayinga GP (Ad26.ZEBOV, Zabdeno) followed by non-replicating modified vaccinia Ankara–vectored vaccine encoding the EBOV Mayinga, SUDV Gulu, and MARV Musoke GPs and the nucleoprotein of the Tai Forest virus (MVA-BN-Filo, Mvabea). The animal rule regulatory pathway, which allows for efficacy data generated in well-characterized animal models of human disease to demonstrate effectiveness in place of phase II or III efficacy trails in humans, facilitated the licensure of the Zabdeno/Mvabea prime-boost vaccination regimen (*13*). This two-shot vaccination is efficacious against SUDV in non-human primates (NHPs) (*14*), but the effectiveness against SVD in humans is unknown.

Furthermore, the two FDA-approved immunotherapeutics, InmazebTM and EbangaTM, target the GP of multiple EBOV strains but do not cross-protect against SUDV (*15*). This lack of protection is expected, given that EBOV and SUDV GPs share less than 56% amino acid identity. A bivalent vaccine that targets both EBOV and SUDV could provide protection against both viruses which have historically caused most outbreaks. Several vaccines that target SUDV (*16*–*20*) or multiple filoviruses (*12*, *21*, *22*) are being developed and tested in clinical trials, including ChAdOx1-biEBOV vaccine.

The ChAdOx1 platform is a replication-deficient simian adenovirus vector, that has been used to generate vaccines which are efficacious against multiple pathogens (*23*–*26*), including severe acute respiratory syndrome coronavirus-2 (SARS-CoV-2). The ChAdOx1 nCoV-19 vaccine against SARS-CoV-2 has been approved in more than 170 countries. ChAdOx1-biEBOV uses the same backbone as ChAdOx1 nCoV-19 but encodes SUDV GP and EBOV GP at the E1 and E4 loci, respectively. ChAdOx1-biEBOV protects type I interferon receptor (Ifnar1[-]) knockout mice against challenge with SUDV Boniface. Here, we evaluated the ChAdOx1-biEBOV efficacy in cynomolgus macaques using either a prime only or prime boost vaccination regimen with SUDV challenge occurring 28 days after the last vaccination. Despite eliciting binding and neutralizing antibodies against SUDV GP, neither vaccine regimen protected against a lethal challenge with SUDV.

## Results

Eighteen female cynomolgus macaques were divided into three groups of six. One group received ChAdOx1-biEBOV 28 days prior to challenge (D-28; prime), one group received ChAdOx1-biEBOV at D-56 and D-28 (prime boost), and one group received ChAdOx1 GFP at D-28 (control). Vaccines were administered via the intramuscular (IM) route with 2.5 x 10^10^ virus particles/animal (Figure 1A). Exams were conducted at the time of vaccination and 14 days thereafter. Serum was collected on days of vaccination, exam, and challenge, and the levels of binding antibodies against SUDV and EBOV GP were measured. Fourteen days post-prime vaccination, antibody levels against SUDV GP varied between 62-1084 arbitrary ELISA units (geometric mean of 294). Fourteen days post-boost vaccination, antibody levels against SUDV GP had increased to 1346-4824 arbitrary ELISA units (geometric mean of 2179). At time of challenge, antibody levels against SUDV GP varied between 30-241 arbitrary ELISA units (geometric mean of 92) for the prime group and 206-882 arbitrary ELISA units (geometric mean of 419) for the prime boost group, 4.6-fold increase over the prime group (Figure 1B). Binding antibodies against EBOV GP followed a similar trend: Antibodies could be detected 14 days post prime vaccination, with arbitrary ELISA units between 50 and 604 (geometric mean of 192) and were increased upon boost vaccination (between 262-1200 arbitrary ELISA units, geometric mean of 618). On the day of challenge, antibodies in the prime boost group were significantly higher (2.3 fold) compared with the prime group (Figure 1C). We investigated the ability of serum antibodies to neutralize SUDV or EBOV using a pseudo-neutralization assay based on rVSV, as previously reported (*17*). Virus-neutralizing (VN) antibodies against SUDV could be detected 28 days after a single vaccination, the VN titers varied between 1:10-1:40. At the time of challenge, all prime-boost sera had a VN titer of 1:40 against SUDV (Figure 1D). Twenty-eight days after a single vaccination, VN titers against EBOV were between 1:10-1:80, whereas all prime-boost sera at time of challenge had VN titers varying between 1:40-1:160 (Figure 1E). The cellular response from peripheral blood mononuclear cells (PBMCs) after vaccination with ChAdOx1-biEBOV was investigated using ELIspot at D-14. Surprisingly, although a robust humoral response was detected, minimal to no interferon-producing T cells were detected upon stimulation with peptides specific for either SUDV GP (Figure 1F) or EBOV GP (Figure 1G) at 14 days post final vaccination.

**Figure 1.**
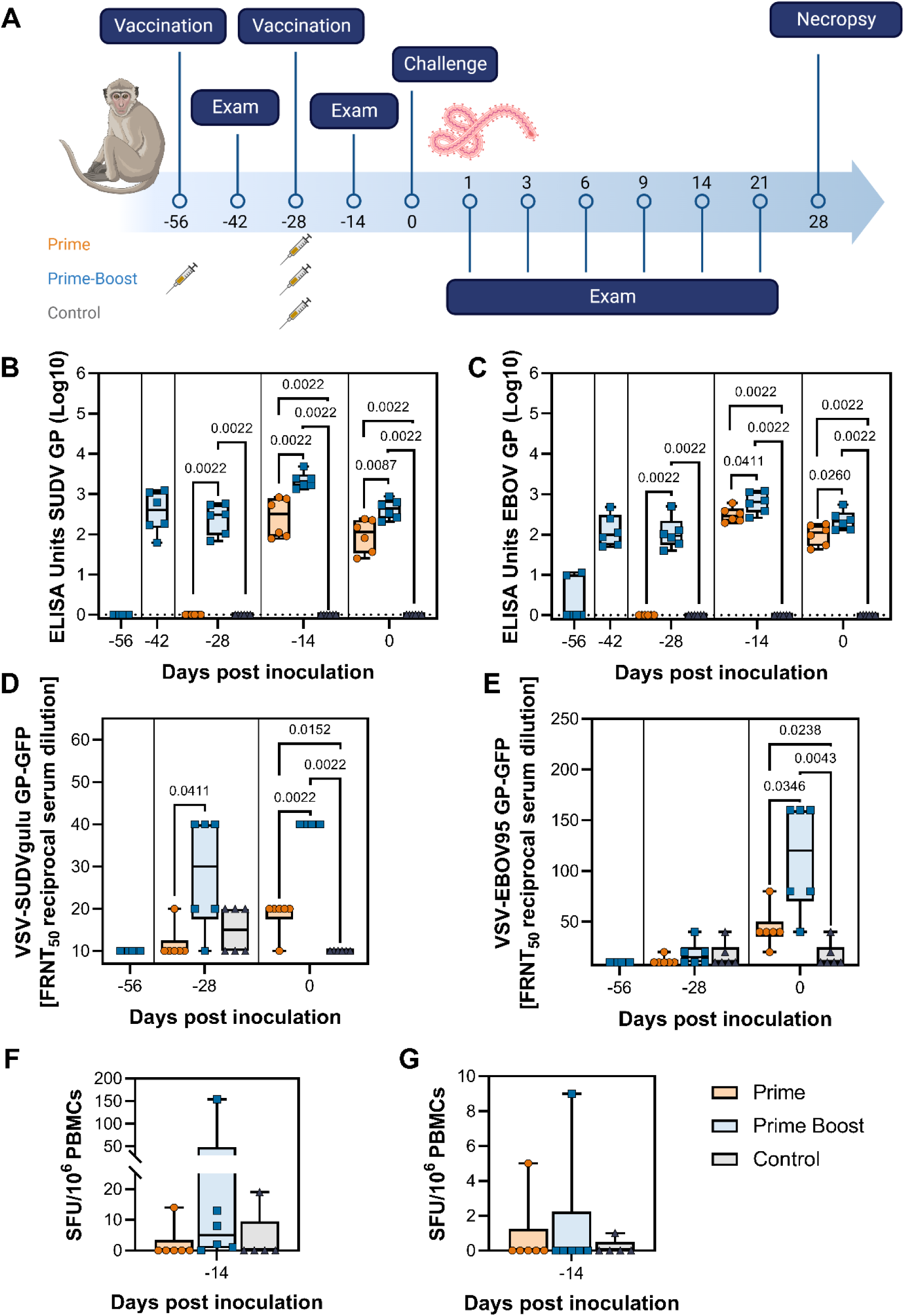
Immunological responses in cynomolgus macaques upon vaccination with ChAdOx1-biEBOV. A. Overview of experimental design. Created with BioRender.com. B/C. Binding antibodies against trimeric SUDV GP (B) or EBOV GP (C) at different time points post vaccination and challenge. Log10 transformed values are shown, which were compared to a standard curve of positive sera. D/E. Virus neutralizing antibodies against pseudovirus VSV-SUDV (D) or VSV-EBOV (E) at different time points post challenge. FRNT50 of green fluorescence protein-positive cell count is shown. F/G. SUDV GP (F) or EBOV GP (G) specific T cell responses in PBMCs isolated from vaccinated or controls animals at D-14 normalized to D−56 or D-28 days post-challenge response. Significance was determined using Kruskall Wallis test followed by Mann Whitney test. The dotted line shows the limit of detection. GP = glycoprotein. FRNT50=50% fluorescence reduction. SUDV=Sudan virus. EBOV=Ebola virus. VSV=vesicular stomatitis virus, SFU=spot-forming units.

Each macaque was challenged with 10^3^ focus-forming units (FFU) SUDV Gulu, administered in 1 mL via the IM route (*17*). Clinical scores for all three groups increased starting on D4, with no significant differences observed between groups (Figure 2A). All macaques developed clinical signs of SVD, including petechial rash in four of six macaques for each group. All were euthanized between D5-7 when they reached end point criteria, and no significant differences in survival were observed between groups (Figure 2B). Viremia was first detected on D3 in two out of six controls and was present in all eighteen macaques on D6, with no significant differences between groups (Figure 2C). Infectious virus was detected in the blood of all 18 macaques at necropsy, with no differences in titers between groups (Figure 2D). Serum chemistry values were significantly different on D6 compared to D0 for the following analytes (Figure 2): increased concentrations of aspartate aminotransferase (AST), increased alkaline phosphatase (ALP), increased blood urea nitrogen (BUN), and decreased albumin. Hematologic values were significantly different on D6 compared to D0 for the following analytes (Figure 2): decreased platelet count, decreased lymphocyte count, and on D3 compared to D0 for increased neutrophil count. Levels of AST and ALP were significantly decreased in the prime boost group compared to the control or prime groups on D6 (Figure 2E-F). Increased blood urea nitrogen (BUN) was observed in all macaques by D6, albeit significantly less in the prime boost group (Figure 2G). Platelet levels in the control group were significantly lower than in the prime boost group on D6 (Figure 2I). Lymphocytes were significantly higher in the prime boost group compared to the control and prime groups on D3 (Figure 2J). Neutrophils were significantly higher in the prime boost group compared to the control group on D3 (Figure 2K).

**Figure 2.**
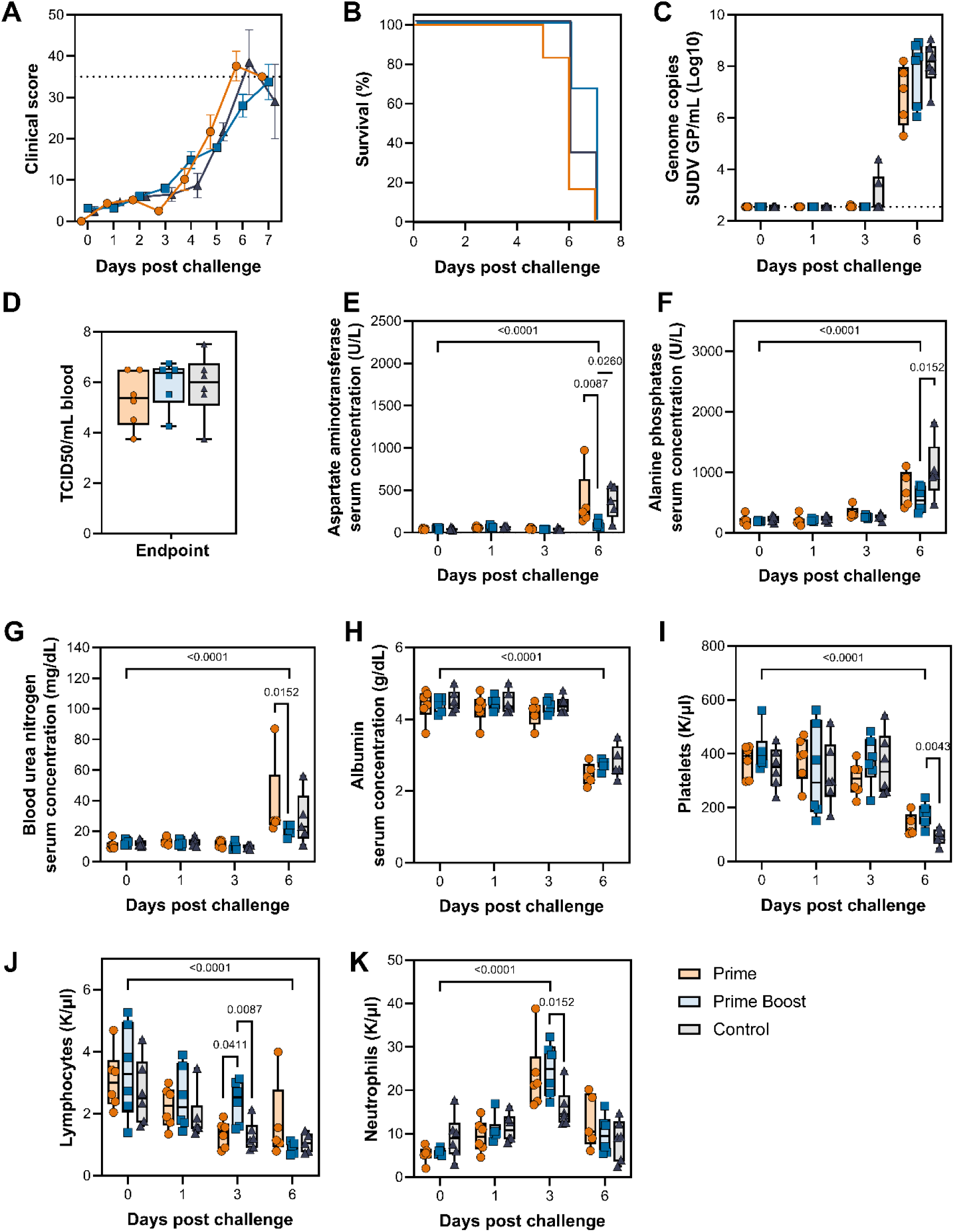
Survival and clinical signs upon challenge of macaques with SUDV Gulu. Clinical score (A) survival (B), and viremia (C) are shown. The following parameters were measured in blood of NHPs: Aspartate aminotransferase (AST, E), alkaline phosphatase (ALP, F), blood urea nitrogen (BUN, G), albumin (H), platelets (I), lymphocytes (J), and neutrophils (K). Significance was determined using Kruskall Wallis test followed by Mann Whitney test. Dotted line shows (A) clinical score at which euthanasia is required, (C) or lower limit of detection.

Although high binding antibody levels against SUDV GP and EBOV GP were detected at challenge, by necropsy the levels of binding antibody had dropped in both vaccine groups, albeit to a lower extent in the prime boost group (Figure 3A-B). We hypothesize that this is due to the presence of sGP and virus with GP on its surface in serum, which would affect our ability to detects SUDV-specific antibodies. Using a SUDV sGP-specific ELISA, sGP was detected in most serum samples obtained on the day of necropsy but not at earlier time points (Figure 3C). Interestingly, we were unable to detect sGP in one sample each for the prime and prime boost group. In these serum samples, SUDV GP-specific binding antibody levels were the highest within the respective vaccine group.

**Figure 3.**
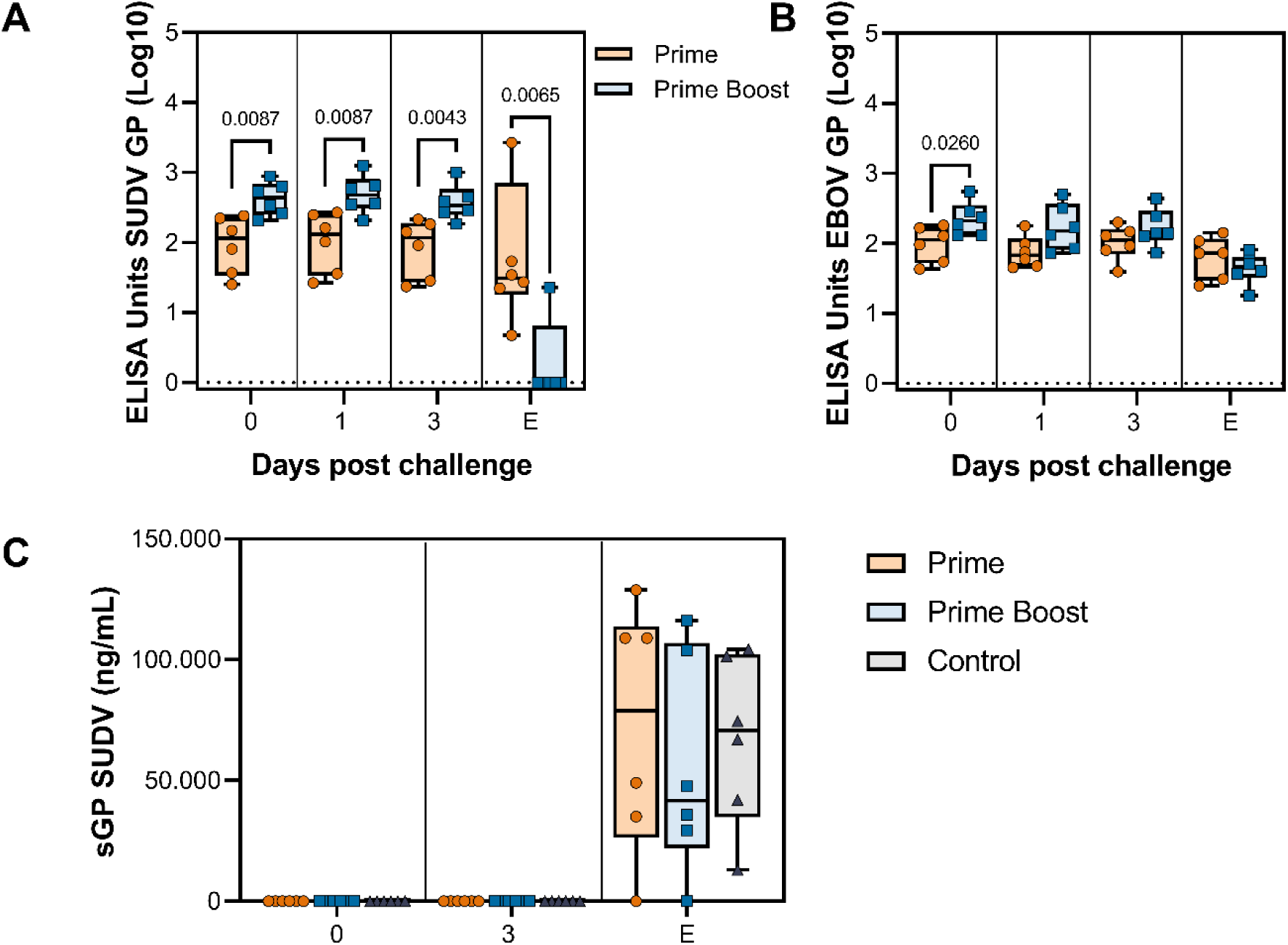
Binding antibody levels and sGP in serum obtained from macaques post challenge. Binding antibody levels against SUDV GP (A) and EBOV GP (B) are shown for the prime and prime boost group post challenge. No binding antibody was detected in control animals. The dotted line shows the limit of detection. C. Soluble GP (sGP) levels were determined in sera using capture ELISA. Significance was determined using Kruskall Wallis test followed by Mann Whitney tests. E = Endpoint. GP = glycoprotein. SUDV=Sudan virus. EBOV=Ebola virus.

A panel of twenty-six different cytokines and chemokines was measured in sera (D-28, 0, 1, 3, and endpoint) as well as in liver and lung tissue homogenates collected at necropsy. In serum (Supplementary Figure 1), an upregulation of multiple cytokines and chemokines could be measured at time of necropsy compared to earlier timepoints (interferon alpha 2 (IFN-α2), interleukin-1 beta (IL-1β), IL-4, IL-6, IL-10, IL-15, IL-16, interferon gamma inducible protein-10 (IP-10), monocyte chemoattractant protein-1 (MCP-1), macrophage inflammatory protein-1 alpha (MIP-1α), MIP-1β, thymus and activation-regulated chemokine (TARC), tumor necrosis factor alpha (TNF-α), and TNF-β). Multiple significant differences were observed between groups at different timepoints, including an increased expression of IFN-ɣ, and decreased expression of IL-4, IL-6, IL-8, IL-15, and IP-10 in the prime boost group compared to the other two groups on D3 or day of necropsy (Supplementary Figure 1).

In liver tissue, multiple cytokines and chemokines were significantly elevated in the prime boost group compared to the prime and control groups (Supplementary Figure 2, IFN-α2, IFN-γ, IP-10, MIP-1α, MIP-1β). No differences in cytokines and chemokines levels were observed between groups in lung tissues.

At time of necropsy, sixteen different samples (liver, spleen, adrenal gland, six lung lobes, inguinal, axillary, mesenteric and mediastinal lymph nodes, aqueous humor, and urine) were collected and investigated for the presence of SUDV viral RNA. Viral RNA was detected in all tissues with no differences between groups detected (Figure 4). The highest viral load was detected in liver, spleen, adrenal gland, and inguinal lymph node. Viral RNA was lowest in aqueous humor.

**Figure 4.**
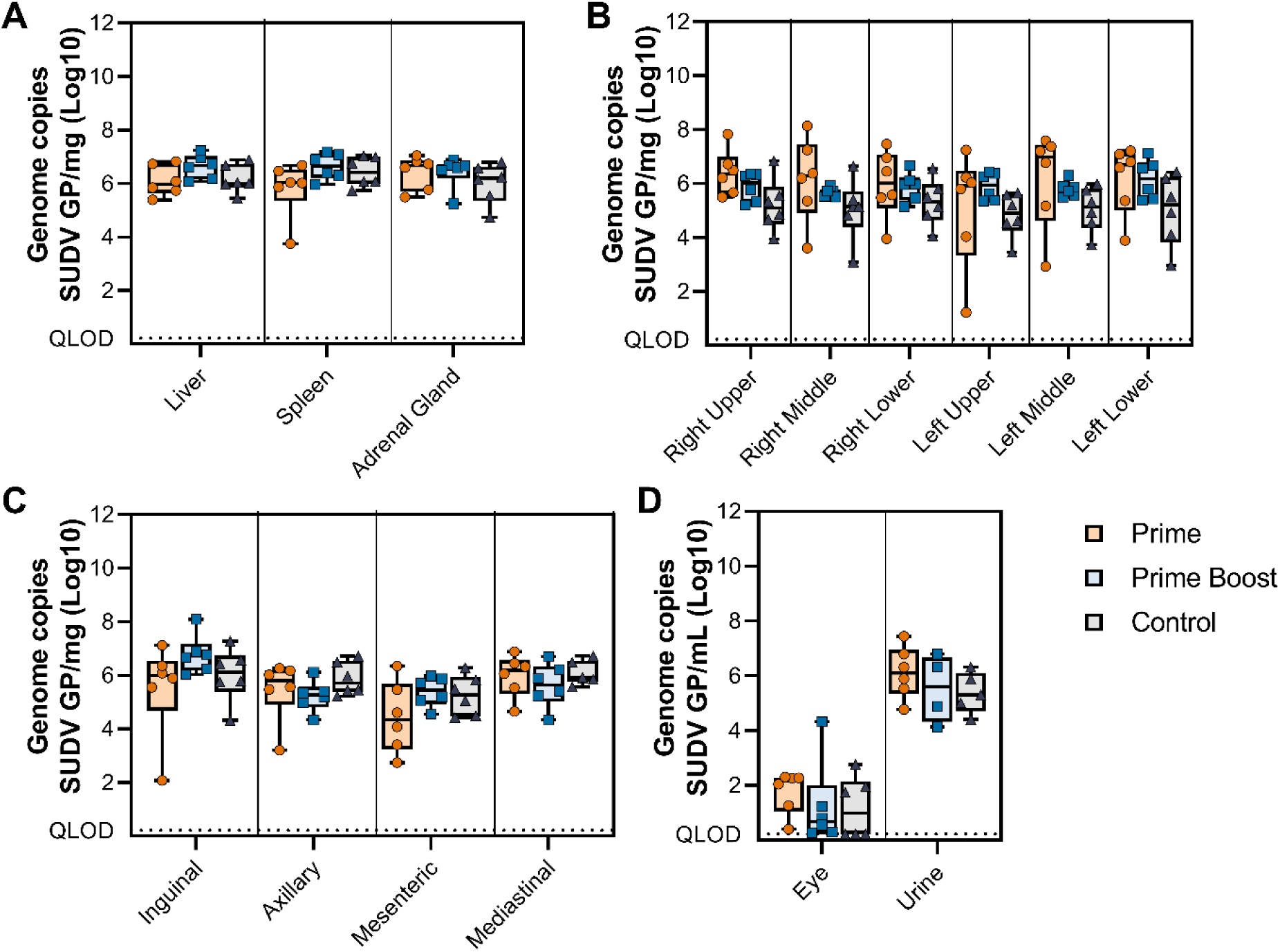
Presence of SUDV viral RNA in tissues obtained from NHPs at necropsy. Viral RNA load was determined in liver, spleen, adrenal gland (A), lung (B), multiple lymph nodes (C), and aqueous humor collected from the eye as well as urine (D). Log10 transformed values are shown, which were compared to a standard curve of known copies of viral SUDV RNA. Significance was determined using Kruskall Wallis test followed by Mann Whitney tests. QLOD = Qualitative limit of detection.

Histologically, all animals demonstrated lesions typical of SUDV infection including multifocal to coalescing hepatocellular degeneration and necrosis with acute inflammation. Splenic lesions consist of abundant micro-fibrin thrombi and splenic white pulp necrosis and loss with abundant fibrin effacing the red pulp. Pulmonary lesions in the control group were minimal and included rare microthrombi in two of six animals and minimal neutrophilic infiltrates in one animal. Prime and prime boost vaccinated animals showed minimal to mild neutrophilic interstitial infiltrates and mild subacute perivascular inflammation in all animals across all pulmonary lobes (Figure 5).

**Figure 5.**
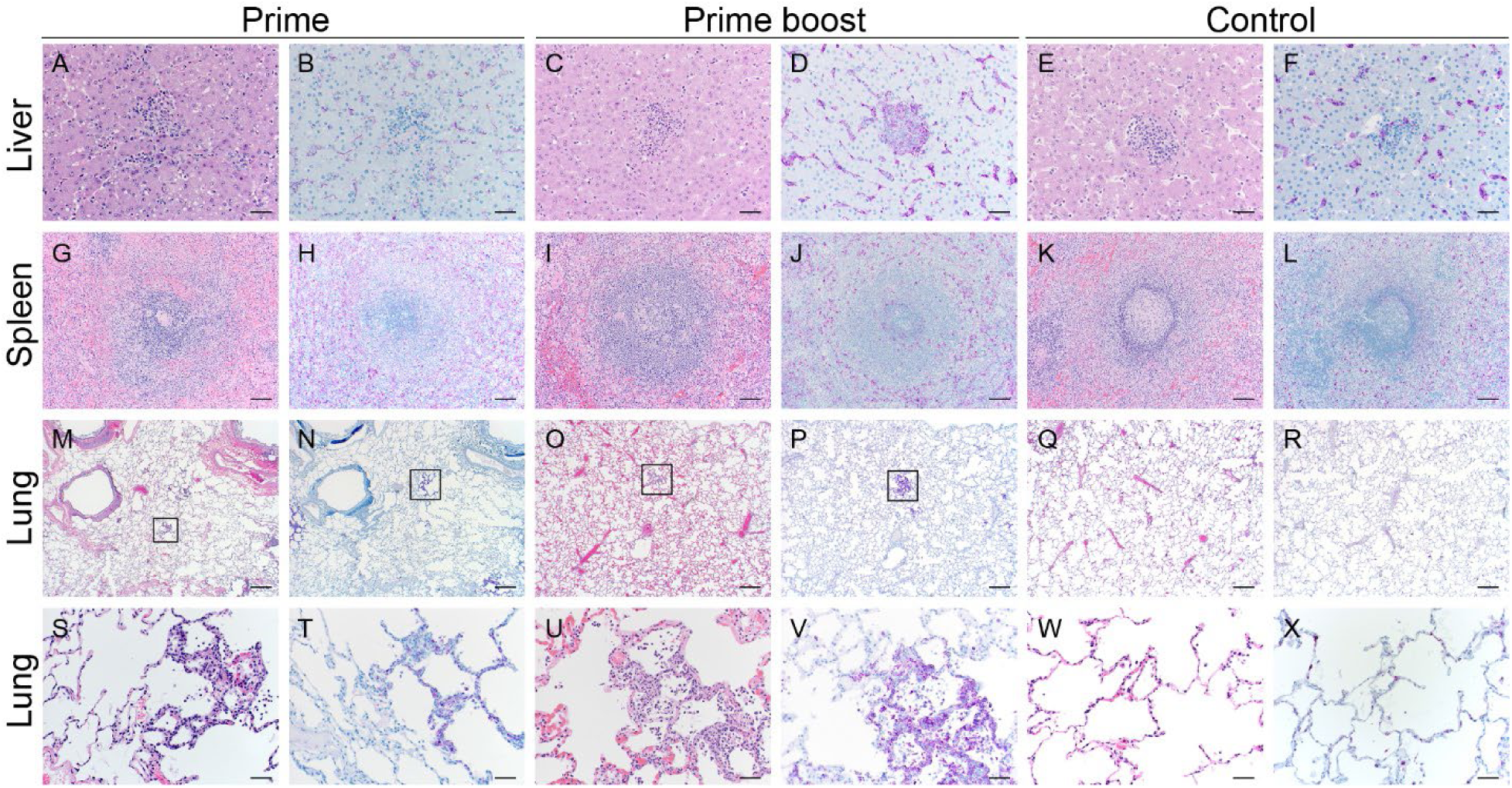
Histopathological effects of ChAdOx1-biEBOV vaccine in the cynomolgus macaque model of SUDV infection. Tissue sections were stained with hematoxylin and eosin (H&E- A,C,E,G,I,K,M,O,Q,S,U,W) or polyclonal rabbit serum against Ebola virus viral protein 40 for detection of viral antigen (purple) by immunohistochemistry (IHC- B,D,F,H,J,L,N,P,R,T,V,X). Liver pathology was similar in all groups consisting of mild-moderate acute inflammation with multifocal hepatocellular necrosis and abundant Kupffer cell and hepatocyte IHC immunoreactivity. Splenic pathology was similar in all groups consisting of minimal-moderate acute inflammation with moderate numbers of red pulp fibrin thrombi; IHC immunoreactivity was prevalent in numerous macrophages. Pulmonary pathology in the prime and prime boost vaccinated animals had minimal-mild neutrophilic interstitial infiltrates shown in the black box (M-P) at 20X magnification and enlarged in S-V at 200X magnification. Control animals did not show evidence of pathology in 5/6 animals with one showing minimal interstitial infiltrates. IHC immunoreactivity in all groups was scatted-numerous in macrophages and endothelial cells. The following magnification was used; liver 200X, spleen 100X, lung 20X and 200X. Bar 200X=50 µm, 100X=100 µm and 20X=500 µm.

## Discussion

We have investigated the efficacy of ChAdOx1-biEBOV against a uniformly lethal challenge with SUDV in cynomolgus macaques. Despite the presence of antigen-specific binding and neutralizing antibodies, we did not observe differences in the clinical signs of disease in vaccinated macaques compared to controls. All macaques reached endpoint criteria and had to be euthanized.

Correlates of protection have not yet been determined for SUDV but have been investigated for EBOV. Recombinant adenovirus serotype 5 (rAd5) encoding EBOV GP is protective against lethal EBOV challenge in cynomolgus macaques. Despite the vaccine eliciting robust EBOV-specific antibodies, passive transfer of purified IgG only protected one out of four macaques. In contrast, CD8+ T cell depletion in vaccinated macaques four days prior to challenge resulted in loss of protection, highlighting the need for T cell mediated responses (*27*). These results parallel with our findings that ChAdOx1-biEBOV immunization resulted in the production of a robust humoral response but did not elicit strong T cell-mediated immunity. Another study utilized immunogenicity and protective efficacy data of the heterologous two-dose vaccine regimen of Ad26.ZEBOV followed by MVA-BN-Filo in a lethal EBOV cynomolgus macaque model and identified binding antibodies against GP as the strongest correlate to survival post-EBOV challenge (*28*). In a previous study using VSV-EBOV vaccination, cynomolgus macaques were T or B cell-depleted before and after vaccination. When they were CD8+ T cell-depleted, they survived a subsequent EBOV challenge four weeks later. CD4+ T cell-dependent survival was time-related: If CD4+ T cells were depleted before vaccination, it reduced EBOV GP-specific antibody development, and macaques did not survive EBOV challenge; whereas, if CD4+ T cells were depleted after vaccination, and macaques were able to develop a normal EBOV-specific antibody response, they survived an EBOV challenge (*29*). Thus, correlates of protection appear to be vaccine platform- and animal model-dependent but suggest an important role for both humoral and cellular immunity dependent on platform.

It should be noted that the virus neutralizing antibody response was higher against EBOV than it was against SUDV using a pseudotype neutralization assay (Figure 1D/E). It is unknown what level of neutralizing antibody is needed to protect against overt disease. A previous VSV-SUDV vaccination study in cynomolgus macaques demonstrated complete protection against a lethal SUDV challenge despite low levels of serum neutralizing antibody titers (*17*). It is therefore likely that a multi-component immune response with high levels of cellular and humoral immunity will be best placed to confer protection in the macaque challenge model.

Without knowledge of correlates of protection against SUDV, which are likely animal model and vaccine platform dependent, NHP studies are helpful to inform selection of vaccine candidates for clinical development. If needed, NHP studies can be performed relatively fast and challenge outcome can be known within five weeks of study start if utilizing a prime only regimen.

Preclinical vaccine studies in NHPs do not necessarily correlate with results obtained in small animal models. Although ChAdOx1-biEBOV was protective against lethal SUDV challenge in Ifnar1[-] mice, this was not the case in cynomolgus macaques. Guinea pigs vaccinated with VSV-EBOV, which provides full protection against guinea pig-adapted (GPA-) EBOV challenge, are partially protected against GPA-SUDV (*30*). However, this finding is not replicated in NHPs (unpublished work). Likewise, we previously showed low immunogenicity of Curevac’s CVnCoV SARS-CoV-2 mRNA vaccine candidate in NHPs (*31*), despite the induction of a humoral response in mice and hamsters and a subsequent reduction in replication of SARS-CoV-2 in hamsters (*32*). Ensuing Phase III clinical trials failed to reach the threshold of 50% vaccine efficacy set by the WHO (*33*). Although small animal models are an essential initial model for vaccine evaluation, when possible, efficacy in NHPs should be evaluated ideally before large-scale clinical trial studies.

Unfortunately, it is not clear from our results why vaccination with ChAdOx1-biEBOV did not provide protection against challenge. Considering that the amino acid identity between SUDV Gulu GP (AAP88031.1), the challenge strain, and Boniface GP (Q66814.1), included in ChAdOx1-biEBOV is 94.67%, we cannot exclude that protective epitopes differ between these strains; however, vaccination with a rVSV expressing SUDV Boniface has conferred protection against SUDV Gulu (*22*). Since the Uganda 2022 outbreak strain’s (OQ672950.1) nucleotide identity is 99.3% with the Gulu strain (KU182912.1) and 94.82% with the Boniface strain (FJ968794.1), SUDV Gulu was selected as the challenge strain. Binding and neutralizing antibodies were elicited against SUDV GP; however, the role of antibodies in protecting cynomolgus macaques against filoviruses when vaccination is done with an adenovirus differs depending on the study design (*27*, *28*). Both binding antibody levels and VN titers were significantly higher in the prime-boost macaques than the prime only, but we did not observe substantial changes in measurements of disease. We measured a minimal T cell response, through ELIspot, after vaccination, even though human volunteers develop a robust cellular response following vaccination with ChAdOx1-biEBOV (unpublished data). We have seen a weak cellular immune response following vaccination of NHPs with the ChAdOx1 vector previously (*23*), in contrast with what has been shown in human volunteers (*34*). This suggests that either the response to ChAdOx1 in NHPs differs from that observed in humans, or that PBMCs are not the correct sample to investigate cellular responses in NHPs vaccinated with ChAdOx1.

As for EBOV, translation of the SUDV GP gene results in three different products: soluble GP (sGP, 70-80%), GP (20-30%), and small sGP (low frequency) (*35*). Furthermore, the presence of sGP reduces neutralizing antibody titer *in vitro* (*36*). Thus, we hypothesized that the presence of SUDV sGP and GP in NHP sera after challenge would be able to bind SUDV GP antibodies and thereby reduce the detection of binding antibodies via ELISA. In line with this hypothesis, the two samples in the prime and prime boost group in which we could not detect SUDV sGP had the highest binding SUDV GP antibody levels at the day of necropsy.

Finally, the EBOV GP-specific response may play a role in the lack of protection against SUDV. *In vitro,* antibodies against EBOV GP have been associated with enhanced cellular entry (*37*). However, it is unclear whether this plays a role *in vivo,* and in this study, no disease enhancement was observed in vaccinees compared to controls. Indeed, when significant differences were observed, they pointed towards a slight improvement of vaccinees over controls, such as in the platelet count. Furthermore, the efficacy of vaccines that combine filovirus antigens in NHPs has yielded variable results depending on the challenge filovirus. Callendret *et al*. mixed Ad26 or Ad35 adenoviral vector-based filovirus GP vaccines expressing Marburg virus (MARV) Angola GP, EBOV Mayinga GP, or SUDV Gulu GP. Cynomolgus macaques were primed with a mixture of vaccines expressing the different GPs in Ad26 vectors, primed with a mixture of vaccines expressing the different GPs in Ad35 vectors four weeks later, and finally challenged four weeks later. Upon lethal challenge with SUDV Gulu, three out of four animals survived (*38*). In a follow-up study with an increased vaccination dose of 4 x 10^10^ virus particles/vector (1.2 x 10^11^ total virus particles/animal), 100% of animals survived a SUDV Gulu challenge, which was significantly correlated to binding antibodies to GP (*14*). A bivalent recombinant protein vaccine that included EBOV Kikwit and SUDV GPs protected all vaccinated cynomolgus macaques from SVD and elicited GP binding antibodies, but the cellular immune response was not evaluated (*39*). A quadrivalent VesiculoVax platform combines four distinct, attenuated rVSVs encoding the GPs of EBOV, SUDV, MARV, and Lassa virus (*22*) or Bundibugyo (*40*) into a single shot. When delivered using a prime-boost regimen, the quadrivalent VesiculoVax protects against SUDV, EBOV, and MARV, elicits production of binding and neutralizing antibodies, and activates antigen-specific T cells in cynomolgus macaques. Of note, IFN-γ producing antigen-specific T cells were only detected in a subset of animals (*22*). When administered a week prior to challenge, all vaccinated animals infected with SUDV, BDBV, or MARV were protected while 80% of macaques challenged with EBOV survived (*40*). Thus, combining different antigens in one vaccine has protected cynomolgus macaques against SUDV in previous studies. A notable difference here is that both antigens were expressed in the same vector; whereas, in the studies described above vaccines encoding a single GP were mixed. When antigens are expressed in the same vector, they will be produced by the same cells, whereas if they are not expressed in the same vector, they may be produced by different cells, potentially resulting in a stronger immune response.

The lack of protection despite vaccine-elicited SUDV GP-specific responses in this model and protection in small animal models suggests that we need a better understanding of the correlates of protection for SUDV vaccines. Overall, this study highlights the importance of evaluating Ebolavirus disease vaccines in multiple models and demonstrates the importance of understanding protective efficacy for any infectious agent in both animal models and human hosts.

## Materials and Methods

### Ethics statement

The Institutional Animal Care and Use Committee of Rocky Mountain Laboratories, National Institutes of Health (NIH) approved all animal experiments. Experiments were carried out in an AAALAC International–accredited facility, following the guidelines and basic principles in the Guide for the Care and Use of Laboratory Animals, the Animal Welfare Act, U.S. Department of Agriculture, and the U.S. Public Health Service Policy on Humane Care and Use of Laboratory Animals. Cynomolgus macaques were singly housed in adjacent primate cages, which allowed for social interactions. The animal room was climate-controlled with a fixed light-dark cycle (12-hour light/12-hour dark). Commercial monkey chow was provided twice daily. The diet was supplemented with treats, vegetables, or fruit daily. Water was available *ad libitum*. Environmental enrichment consisted of a variety of human interaction, manipulanda, commercial toys, videos, and music. Animals were monitored at least twice daily. The Institutional Biosafety Committee approved work with SUDV under biosafety level 4 conditions and subsequent sample inactivation for removal of specimens from high containment (*41*, *42*).

### Generation of ChAdOx1-biEBOV

Generation of ChAdOx1-biEBOV was done as described previously (Flaxman GP paper if published). Briefly, EBOV GP (Makona-Kissidougou-C15 GenBank:KJ660346.1) and SUDV GP (Boniface, UniProtKB Q66814.1) were codon-optimized for humans. Antigen cassettes were cloned into an ENTRY plasmid and recombined *in vitro* with ChAdOx1-biDEST containing a Gateway destination cassette in both the E1 and the E4 loci, respectively. E1 and E4 expression cassettes both contained the TetR-repressible CMV promoter; E1 cassettes contained the BGH polyA sequence, while E4 expression cassettes contained the SV40 polyA sequence. SUDV GP was encoded in the E1 cassette, whereas EBOV GP was encoded in the E4 cassette. Viral vectors were produced at the Viral Vector Core Facility at the Jenner Institute. All adenovirus vectors were produced in the T-REx-293 cell line (Thermo Fisher Scientific), which allows for transcriptional repression of the vaccine antigens during vector production. Vectors underwent quality control (including titration, identity PCR, and sterility testing) before being used in *in vitro* and *in vivo* studies.

### Study design

Eighteen cynomolgus macaques of Indonesian origin, all female, were sorted by weight and then randomly divided into three groups of six animals each (prime group 8-11 years old, 3.2-6.0kg; prime boost group 8-13 years old, 3.2-5.2kg; control group 7-11 years old, 3.4-4.7kg). Animals were vaccinated with 2.5 x 10^10^ virus particles per animal of ChAdOx1-biEBOV or ChAdOx1 GFP intramuscularly (250 µl in each caudal thigh), as done previously (*23*). Vaccine was diluted in sterile phosphate-buffered saline (PBS). All animals received a vaccination at D-28, and the prime boost group received an additional vaccination at D-56. On D0, animals received 1 x 10^4^ TCID_50_ of SUDV Gulu (500 µl in each caudal thigh). Clinical scoring was done daily post challenge and based on the evaluation of the following criteria: general appearance and activity, appearance of skin and coat, discharge, respiration, feces and urine output, and appetite. Clinical exams were conducted on D−56, −42, −28, −14, 0, 1, 3, and 6. All animals reached endpoint criteria between D5-7. Hematology analysis was completed on a ProCyte DX (IDEXX Laboratories, Westbrook, ME, USA) and the following parameters were evaluated: red blood cells (RBC), hemoglobin (Hb), hematocrit (HCT), mean corpuscular volume (MCV), mean corpuscular hemoglobin (MCH), mean corpuscular hemoglobin concentration (MCHC), red cell distribution weight (RDW), platelets, mean platelet volume (MPV), white blood cells (WBC), neutrophil count (abs and %), lymphocyte count (abs and %), monocyte count (abs and %), eosinophil count (abs and %), and basophil count (abs and %). Serum chemistries were completed on a VetScan VS2 Chemistry Analyzer (Abaxis, Union City, CA) and the following parameters were evaluated: glucose, blood urea nitrogen (BUN), creatinine, calcium, albumin, total protein, alanine aminotransferase (ALT), aspartate aminotransferase (AST), alkaline phosphatase (ALP), total bilirubin, globulin, sodium, potassium, chloride, and total carbon dioxide. Upon reaching endpoint criteria, animals were euthanized, and necropsy was performed. The following tissues were collected: muscle at injection site, liver, spleen, adrenal gland, 6 lung lobes, inguinal, axillary, mesenteric and mediastinal lymph nodes, aqueous humor, and urine. Urine was not collected from three of the macaques, one in each study group.

### Cells and Virus

SUDV Gulu (NC_006432.1) was obtained from the US Army Medical Research Institute of Infectious Diseases and propagated once on Vero E6 cells. Deep sequencing revealed no contaminants; however, compared to the reference sequence, four base pair changes (two in NP, one in VP40, and one non-coding in GP (*17*)) were noted. VeroE6 cells were maintained in DMEM supplemented with 10% fetal bovine serum, 1 mM l-glutamine, penicillin (50 U/ml), and streptomycin (50 μg/ml; DMEM10). Mycoplasma testing is performed at regular intervals, and no mycoplasma was detected.

### RNA extraction and quantitative reverse-transcription polymerase chain reaction

RNA was extracted from whole blood using a QiaAmp Viral RNA kit (Qiagen) according to the manufacturer’s instructions. Tissue was homogenized and extracted using the RNeasy kit (Qiagen) according to the manufacturer’s instructions. A SUDV GP viral RNA–specific assay (F: CAAAGGGAAGAATCTCCGACC, R: CAGGGGAATTCTTTGGAACC, P: FAM-GGCCACCAGGAAGTATTCGGACC-BHQ; designed using the Sudan ebolavirus isolate Ebola virus/H.sapiens-tc/UGA/2000/Gulu-808892 (KR063670.1) sequence) was used for the detection of viral RNA. RNA (5 μl) was tested with QuantStudio (Thermo Fisher Scientific) according to instructions of the manufacturer. To create standards, SUDV Gulu stocks were extracted using QIAamp viral RNA kits per manufacturer’s instructions. RNA was serially diluted and quantified using the One-Step RT-ddPCR system (Bio-Rad). Briefly, 5 µL of RNA was combined with recommended volumes and concentrations of supermix, reverse transcriptase, DTT, primers, probe, and molecular grade water. This mix was then combined with Automated Droplet Generation Oil for Probes using the Automated Droplet Generator. Once the droplets were generated, PCR was performed using recommended cycling conditions. Reverse transcription was performed at 50°C and anneal/extension was performed at 60°C. After thermal cycling, positive droplets were quantified with the QX200 Droplet Reader. Finally, a range of copies/rxn that gave cycle threshold values between 20-30 were selected as controls and used to quantify RNA in this study.

### SUDV GP and EBOV GP ELISA

The presence of SUDV GP- and EBOV GP-specific binding antibodies in macaque sera was determined using ELISA, carried out in Nunc MaxiSorp flat bottom 96-well plates (Thermo Fisher Scientific) coated with 50 ng/well of trimeric ectodomain SUDV GP or EBOV GP (NativeAntigen) in DPBS followed by overnight incubation at 4°C. The wells were blocked with 100 µl of casein in PBS (ThermoFisher) for 1 h at room temperature and then incubated with diluted sera in triplicate for 1 h at 37°C. Serially diluted (1:100 to 1:51200) sera obtained from a SUDV-challenged macaques at 40 days post challenge was utilized as a standard curve and diluted 1:1600 as an internal control. IgG antibodies were detected by using affinity-purified polyclonal antibody peroxidase labelled goat-anti-monkey IgG (Seracare) at a dilution of 1:2500 in casein in PBS for 1 h at 37°C. The absorbance was measured at 405 nm after the addition of TMB two component peroxidase substrate (Seracare, 5120-0047) for 5-10 min followed by the addition of stop solution (Seracare, 5150-0021). Wells were washed 6x with PBST 0.1% Tween in between each step. Arbitrary ELISA units were calculated based on the standard curve. All samples diluted at 200x with an OD value of less than 0.250 were given a value of 1.

### sGP ELISA

The SUDV sGP-specific sandwich ELISA was performed as previously described (*43*) with minor differences. SUDV sGP was captured using the 1 µg/mL polyclonal rabbit anti-SUDV sGP antibody (cat. # 0302-030, IBT Bioservices). Dilutions of recombinant SUDV sGP (cat. # 0570-001, IBT Bioservices) at known concentrations served as standards.

### Pseudovirus neutralization

Neutralization assays with VSV-EBOV-GFP and VSV-SUDV-GFP were performed as previously described (*17*). The VSV-SUDV-GFP assay was optimized to include incubation of serum dilution mix on the Vero E6 cells for 16 hours at 37°C. Samples were analyzed on the FACSymphony^TM^ A5 Cell Analyzer (BD Biosciences, San Jose, CA, USA) and the GFP-positive cell count was analyzed using FlowJo v10 Software (BD Life Sciences).

### ELIspot assay

Peripheral blood mononuclear cells (PBMCs) were isolated from ethylene diamine tetraaceticacid (EDTA) whole blood on certain exam days (D-56, -42, -28, -14, and 0) using Leucosep tubes (Greiner Bio-one International) and Histopaque-1077 density-gradient cell-separation medium (Sigma-Aldrich) according to the manufacturers’ instructions and frozen in liquid nitrogen. After thawing of cells, cells were plated at a concentration of approximately 100,000 cells per well and were stimulated for 6 hours with contiguous peptide pools that spanned the length of the EBOV or SUDV GP at a concentration of 2 μg/mL per peptide. As a positive control, cells were stimulated with eBioscience™ Cell Stimulation Cocktail (ThermoFisher). As a negative control, cells were incubated in media with equivalent amounts of DMSO. The ImmunoSpot Human IFNγ Single-Colour Enzymatic ELISpot Assay Kit was performed according to the manufacturer’s protocol (Cellular Technology). Analysis was performed using the CTL ImmunoSpot Analyzer and ImmunoSpot Software (Cellular Technology).

### Measurement of cytokines and chemokines

Sera was collected on all exam days. Tissue (Lower left lung lobe, and liver) was weighted, homogenized in 750 µl of DMEM, spun down for 10 min at 8000 rpm, after which supernatant was collected. Sera and tissue supernatant were irradiated (4 MRad) and analyzed on the MesoQuick Plex (MSD, K15203D) at a 4x and 10x dilution, respectively. The U-PLEX Biomarker Group 1 (NHP) Assay kit (MSD, K15068L) was used to test the presence of twelve cytokines (GM-CSF, IFNα2a, IFN-γ, IL-1β, IL-4, IL-6, IL-10, IL-12 p70, IL-15, IP-10, MCP-1, and TNF-α), the V-PLEX Cytokine Panel 1 Human kit (MSD, K15050D) was used to test the presence of seven additional cytokines (IL-5, IL-7, IL-12/IL-23p40, IL-16, IL-17A, TNF-β, and VEGF-A), and the V-PLEX Chemokine Panel 1 Human kit (MSD, K15047D) was used to test the presence of seven additional chemokines (Eotaxin-3, IL-8 (HA), MCP-1, MDC, MIP-1α, MIP-1β, and TARC). Data were extracted from the plates using the MSD Workbench 4.0 software and qualitative lower limit of detection (QLOD) were calculated for each parameter. All values below the QLOD were changed to the QLOD. Cytokine and chemokine concentrations were calculated per mg of tissue and mL of sera.

### Histology and immunohistochemistry

Necropsies and tissue sampling were performed according to IBC-approved protocols. Tissues were perfused with 10% neutral-buffered formalin and fixed for a minimum of 7 days. Tissues were placed in cassettes and processed with a Sakura VIP-6 Tissue Tek, on a 12-hour automated schedule, using a graded series of ethanol, xylene, and PureAffin. Embedded tissues are sectioned at 5 µm and dried overnight at 42°C prior to staining. Specific anti-Ebolavirus immunoreactivity was detected using an in-house rabbit polyclonal anti-Ebola Zaire VP40 antibody at a 1:2000 dilution. Vector Laboratories ImmPress VR horse anti-rabbit IgG polymer (# MP-6401) was used as a secondary antibody. The tissues were stained using the Discovery Ultra automated stainer (Ventana Medical Systems) with a Roche Tissue Diagnostics Discovery purple kit (#760-229). The tissue slides were examined by a board-certified veterinary anatomic pathologist blinded to study group allocations. Pulmonary pathology was evaluated in three histological sections from each of the six lung lobes resulting in eighteen evaluated sections for each macaque. Histologic lesion severity was scored according to a standardized scoring system evaluating the presence of pathology: 0, no lesions; 1, minimal (1-10% of lobe affected); 2, mild (11-25%); 3, moderate (26-50%); 4, marked (51-75%); 5, severe (76-100%). Presence of viral antigen was scored per lung lobe according to a standardized scoring system: 0, none; 1, rare/few; 2, scattered; 3, moderate; 4, numerous; 5, diffuse. A representative lesion from each group was selected for figures.

### Statistical analysis

Kruskall-Wallis analysis was conducted to compare differences between groups, followed by Mann-Whitney test if differences were found to be significant using GraphPad Prism version 8.3.0. Statistical tests used are identified in the figure legends.

## Author contributions

SvT, AM, TL, VJM, and NvD designed the study; AM, TL and VJM acquired funding; SvT, PF, FF, RKM, JRP, SG, JES, JR, RM, AC, LM, JL, BJS, AO, CS, GS, AM, NvD acquired, analyzed, and interpreted the data; SvT and NvD wrote the manuscript. All authors have approved the submitted version.

## Acknowledgements

We thank Brandon Bailes, Viviane Callier, Lydia Crawford, Richard Cole, Kathy Cordova, Allison Darrow, Dean Follmann, Corey Henderson, Taylor Lippincott, Kay Menk, Kyle O’Donnell, and Taylor Saturday for their assistance during the study.

---

**Supplementary Figure 1.**
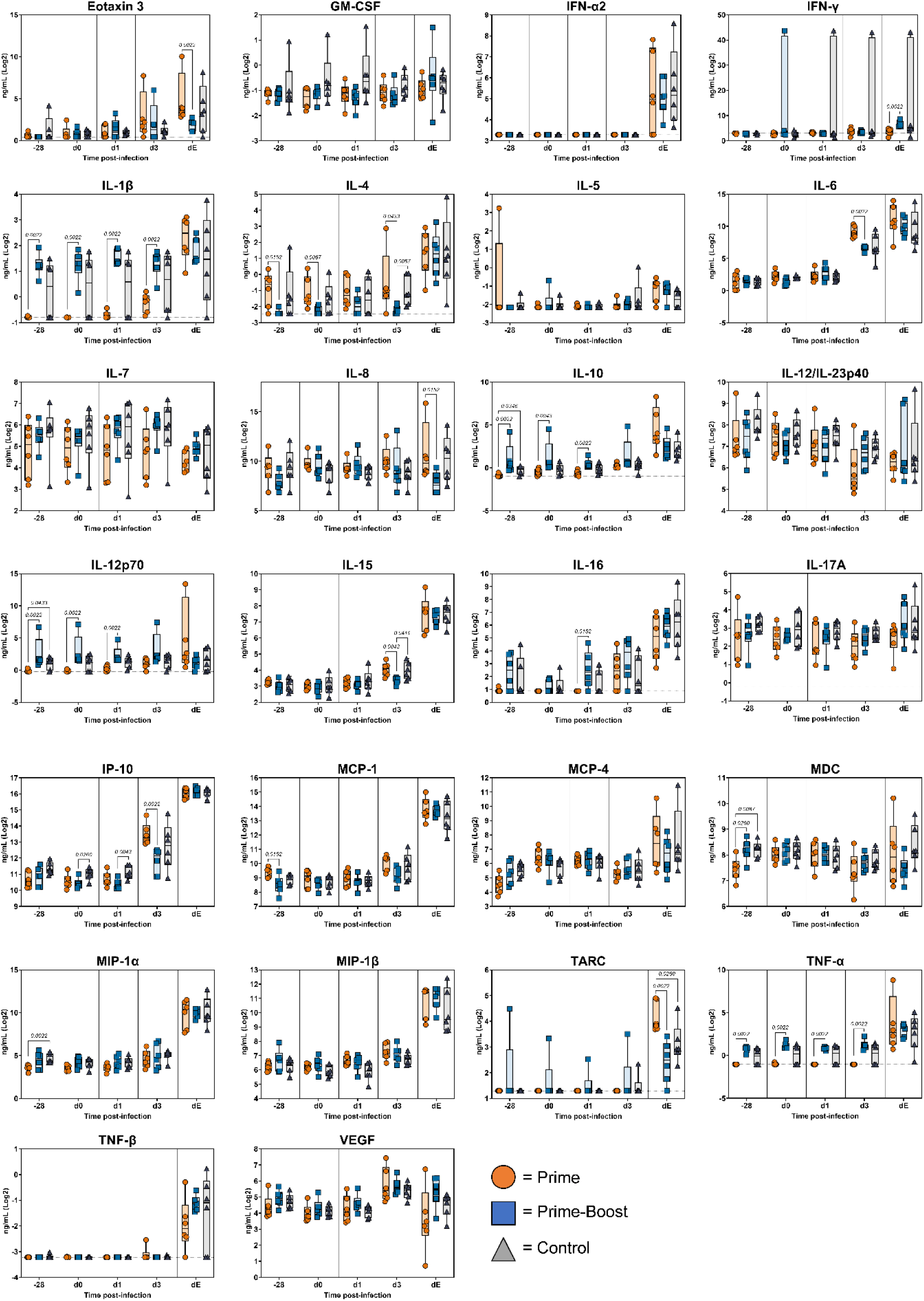
Cytokines and chemokines in serum from cynomolgus macaques challenged with SUDV. Serum was collected at exam and cytokine and chemokine concentrations were measured using the Meso Quickplex.

**Supplementary Figure 2.**
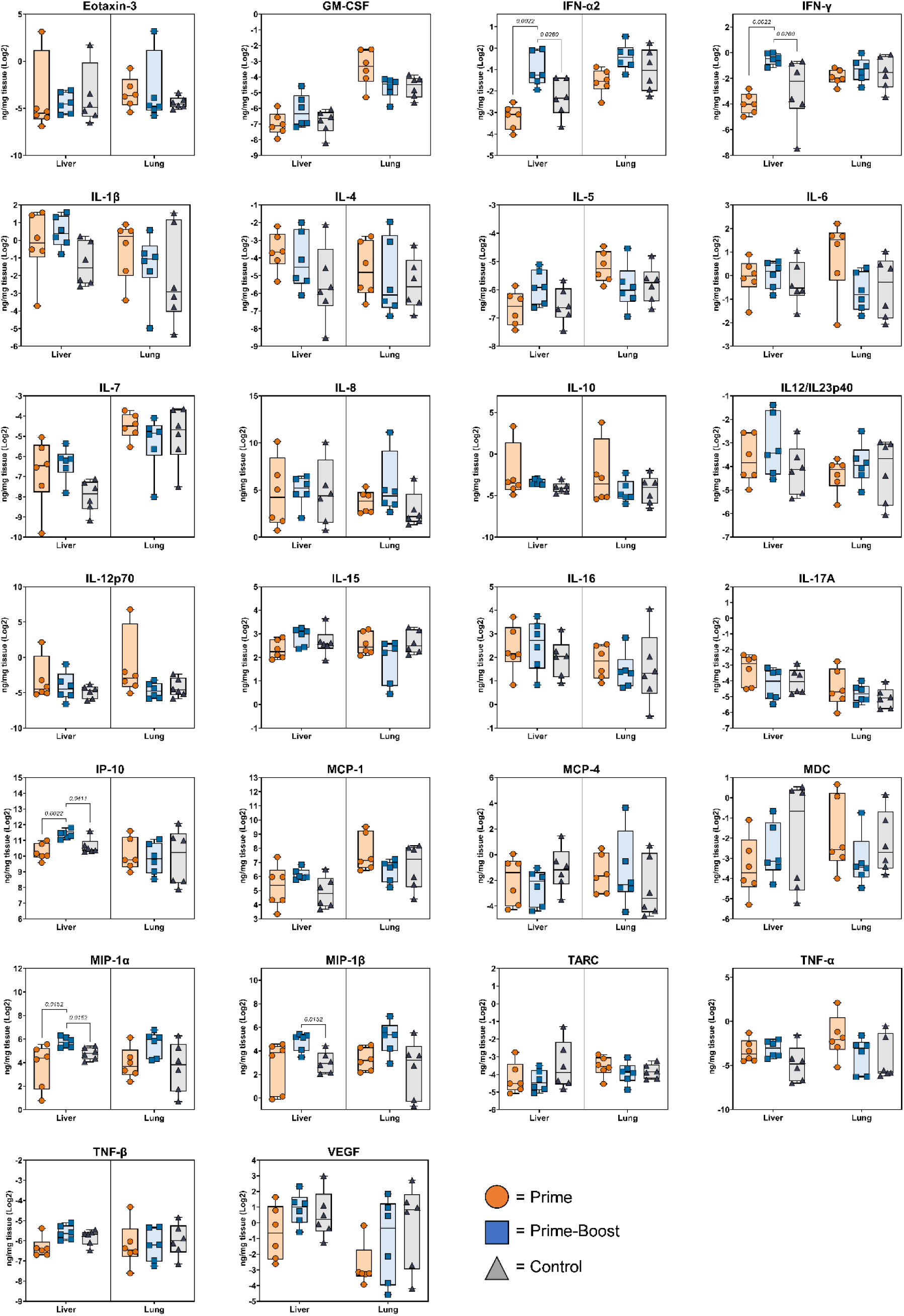
Cytokines and chemokines in tissues from cynomolgus macaques challenged with SUDV. Tissues were collected at necropsy and cytokine and chemokine concentrations were measured using the Meso Quickplex.

